# Ecological and spatial overlap indicate interspecific competition during North American Canid radiation

**DOI:** 10.1101/2023.12.14.571772

**Authors:** Rodolfo P. Graciotti, Lucas Porto, Tiago B. Quental

**Affiliations:** Universidade de Sao Paulo

## Abstract

Understanding biodiversity patterns and the processes that generate them are key goals in ecology and macroevolutionary studies. Diversity-dependent models of diversification have been used to indirectly infer the relevance of interspecific competition on driving speciation and extinction dynamics. In this study, we develop a new approach that more explicitly incorporated spatial and eco-morphological overlap among species to test how interspecific competition my affect diversification dynamics at deep time. We built different metrics that capture not only species temporal coexistence, but also their coexistence in space and morphospace to test the hypothesis that an increase in the intensity of competition would result in both a decrease in speciation rate and an increase in extinction rate. We tested our predictions using the North American fossil record of the family Canidae, a group that has been extensively studied and well characterized both from the eco-morphological and paleontological points of view. We found that interspecific competition only affected diversification dynamics during the early stages of Canidae radiation, resulting only in the suppression of speciation rate at the time the clade was expanding in diversity. We found no association between the intensity of the competition and extinction dynamics as expected by a competitive model, nor an association between changes in speciation and extinction rates and changes in global temperature, suggesting that extinction dynamics might be more related to external factors, such as clade-competition. We discuss the relevance of different factors on driving diversification dynamics changes over time and how evaluating the role of interspecific competition using different metrics that better capture the intensity of competition (as opposed to diversity dependent models), might be a way forward to investigate the role of biotic interactions at deep time.

## Introduction

Macroevolutionary studies are often directed towards understanding how species diversity changed in deep time, a multimillion-year time scale. The ultimate goal of such studies is to reveal the underlying processes that shape those patterns (Jablonski, 2008; Rabosky 2013; Hembry & Weber 2020). Traditional views on the controls of diversity through time often antagonize the roles of the abiotic environment and the biotic interactions between species (Barnosky, 2001), however those different factors are not self-excluding. Biotic and abiotic factors could operate at different temporal scales (Benton, 2009), at different moments during the radiation of a given clade (Neubauer et al 2022) or interact to simultaneously affect diversification dynamics (Ezard et al., 2011). Although abiotic factors have captivated the interest of most paleontologists, the recent advent of new approaches (Fraser et al 2018) has revitalized the interest on biotic factors as an important regulator of biodiversity at deep time.

Previous palaeobiological studies have inferred the role of biotic interactions on diversification rate dynamics in a very indirect manner. Those have focused on the role of competition by either showing that the diversification dynamics of different lineages are coupled (e.g. Liow at all 2015), or found evidence in favor of diversity-dependent diversification (e.g. Naubeuar et al 2023; Pires at al 2017). Studies on carnivorous mammals have been particularly important to suggest an active role for competition in driving diversification dynamics (Silvestro et al., 2015; Van Valkenburgh, 1988, 1999). Van Valkenburgh (1988, 1996) has long suggested that competition might operate among clades, and Silvestro et al (2015) has shown evidence for both within and among clade competition affecting the diversification rate dynamics *per se*. More recently, Pires et al. (2017) has proposed that speciation dynamics may be driven by competitive interactions between species within a given clade while the interaction between species in different lineages (clade competition) more often results in changes in extinction regimes.

The idea that competitive interactions may be a driving force in speciation or extinction is central to the theory of diversity-dependent diversification (Phillimmore and Price, 2008; Rabosky, 2013;but see Moen & Morlon 2014) but we should note that this link is a rather indirect evidence of competition. Under this tenet, diversity-dependence has been interpreted as being evidence of competition between species that overlap in space and time (Sepkoski, 1978, 1996), through mechanisms such as competitive exclusion acting on extinction, or limited resources acting on suppressing speciation (Rabosky, 2009, 2013; Sepkoski, 1978, 1996). Such competition could happen either within species of a given clade or among species of distantly related clades (Krause 1986; Sepkoski 1996; Lupia et al 1999; Van Valkenburgh 1999; Liow at all 2015; Silvestro et al 2015; Pires et al 2017). It is important to note that most previous studies consider ecological and spatial overlap in a very crude sense, either by cherry picking lineages that are likely to interact, or by restricting the analysis to regional, rather than global scale. In this study, we developed an approach that more explicitly incorporates ecological mechanisms. More specifically, we develop a metric to describe the intensity of interspecific competition that explicitly takes into account ecological and spatial overlap. We used the fossil record of north American canids to ask the question whether interspecific competition influences both the speciation and extinction regimes and at what stages of the radiation is interspecific competition relevant, if at all.

## Methods

### Studied organism

The family Canidae (Mammalia) comprises both extant and extinct lineages. It originated in the Paleogene of North America, having existed for some 40 million years until the present day (Wang & Tedford, 2008). The Canidae fossil record of North America is considerably well documented, not only taxonomically, but it shows a comparatively high level of completion when compared to other vertebrates (Tedford et al., 2009; Wang, 1994; Wang et al., 1999). Extinct species have also been well characterized eco-morphologically (Balisi et al., 2018; Balisi & Van Valkenburgh, 2020; Janis et al., 1998; Slater, 2015; Van Valkenburgh et al., 2004). Extinct canid species vary considerably in both body size and diet, ranging from small predators of small prey and plant matter (defined as hypocarnivory), to large-bodied bone-crushing or hypercarnivore dogs, whose diet is almost entirely composed of meat (Van Valkenburgh, 1991; Wang & Tedford, 2008; Slater, 2015).

### Fossil occurrence and ecomorphological data

We retrieved fossil occurrence data from Paleobiology Database (PBDB), restricting our analyses to include only North American canids, as most of the clade evolutionary history was confined in this continent (Wang & Tedford, 2008). Curatorial work on the data followed Pires et al. (2015), and the taxonomic treatments of several different authors (Tedford et al., 2009; Wang, 1994; Wang et al., 1999; Wang & Tedford, 2008, Zrzavý et al., 2018). All occurrences with a temporal range estimated to be larger than 15 million years were removed (Figure S1). Our original dataset comprised 2210 occurrences and after our successive curatorial work, we ended up with 1555 occurrences of 138 species. Each occurrence is georeferenced with latitude and longitude coordinates, and that information was used to estimate spatial overlap. Please see the supplemental material for details on the curatorial work done for the occurrence data.

To categorize their eco-morphology we described each species in a two-dimensional morphospace that encompasses two important morphological proxies for species niche: body size and a metric that is a proxy for diet (the level of carnivory). We used Balisi & Valkenburgh (2020) published dataset of canid body mass (see supplemental methods), with data for 125 species out of 138 species for which we had fossil occurrence data. The original dataset described body mass in kilogram units, but we took the natural logarithm before running our analysis. To infer diet, we used the results of a Linear Discriminant Analysis (LDA) from Slater (2015) based on several craniodental variables (see supplemental methods for details). This analysis classifies each Canidae species in discrete diet categories of hypercarnivory, mesocarnivory and hypocarnivory. We used both this discrete category, as well as the first axis of LDA (LD1) as a continuous measure of carnivory. This allowed us to classify 97 out of 138 species with occurrence data. We designed a data augmentation-approach to complete the morphology of species we had no eco-morphological data (see supplemental material for details). We note that this data augmentation should not generate any bias given that it was a taxonomically oriented random data input procedure replicated several times to incorporate its uncertainty. We repeated the whole data augmentation process several times ending up with 500 different morphological datasets. If anything, the procedure introduces white noise and hence could be conservative with respect to finding significant patterns (please see the supplemental methods for details on how this was done).

### Analyses

We proceed our analyses in a three-step framework: first, we analyzed fossil occurrence data in a Bayesian framework to estimate the “true” times of origin and extinction for each species, the preservation and diversification rates. Second, we implemented a modular framework to describe different aspects of species coexistence and the potential for competition. This was done by implementing different metrics of disparity and measuring spatial overlap. The scenarios considered included those that do not explicitly take into account spatial and morphological overlap (a simple diversity dependence), scenarios that take into account either spatial (diversity dependence at different spatial scales) or morphological overlap, a scenario that take into account both spatial and ecological aspects. Each different scenario produced a time series that described how the intensity of competition varied through time. The last step consisted of using these time series to investigate the potential effects of biotic interactions on macroevolutionary dynamics. By incorporating the changes observed in competition dynamics through the time series in a Bayesian framework, we were able to investigate if such changes were associated with changes in speciation and extinction rates.

### Initial PyRate analysis

We analyzed fossil occurrences in *PyRate*, a hierarchical Bayesian framework that jointly estimates preservation and diversification processes, while explicitly incorporating different aspects of the incompleteness of the fossil record, using all known occurrences of a given lineage (Silvestro et al., 2014, 2019). *PyRate* works on a fully probabilistic Bayesian paradigm, allowing us to quantify the levels of uncertainty of such estimates. We utilized all our occurrences of Canidae North American fossil record to estimate species times of speciation and extinction, and speciation and extinction rates. We set our analyses using the mG + qShfit model, running 30000000 RJBDMCMC iterations, sampling every 10000 iterations to obtain the posterior estimates of the parameters, discarding the first 10% as *burn-in*. Those settings allowed us to model a preservation processes that simultaneously accounts for preservation heterogeneity both through-time (using a Time-variable Poisson Process - TPP) and across lineages, an approach deemed more realistic to describe varying patterns of fossil preservation (the mG + qShift model in *PyRate* notation). Because most of Canidae fossil occurrences are associated to the North American Land Mammal Ages (NALMAs), we grouped some of these stages to define the time windows where preservation could vary: we set the time window boundaries at 37.2, 30.8, 20.43, 15.97, 13.6, 10.3, 4.9, 1.8, 0.3 and 0 Million years ago (Ma). We utilized such times frames to accommodate a roughly homogeneous number of occurrences among intervals across the whole-time frame (Figure S2). We set the number of living species to be 9 given that this is the number of Canids seem today in North America. To account for the uncertainty associated with the age of each fossil occurrence, we randomly drew ages within each occurrence timespan, generating 50 different temporal replicates and conducted *PyRate* analyses on each of these replicated datasets. We restricted fossil occurrences found in the same assemblage to be assigned the exact same age, as they may represent the same fossilizing event. Please see the supplemental material for further details. From these analyses we described the general trend in speciation and extinction rates, as well as the times of origin and extinction of each species, and hence their longevities. This information was then used as inputs in the step used to define coexistence in time described in the next sections.

### Modular framework

In our attempt to better describe competition dynamics, we developed a modular approach to capture different levels of complexity of species interaction in the fossil record. We designed and explored four different scenarios that could be used as proxies for detecting competition as follows:

a. competition intensity is described by the variation in the absolute number of species coexisting through time, a classic diversity dependence original scenario, where all species that coexist in time at the North American continent are considered potential competitors, irrespective of their ecology or local spatial distribution.
b. competition intensity is described by the variation in the number of species co-occurring in time and at different spatial scales. To be considered a potential competitors different species have to spatially co-occur at the local spatial scale (see details below).
c. competition is described by how disparity metrics of morphospace density change through time, without considering if they coexist in space. Here, all species coexisting through time are potential competitors and competition intensity is measured by species distance in the morphospace.
d. competition is described by how the disparity metrics of morphospace density change through time, modulated by temporal and spatial coexistence (at different scales), hence not all species are considered potential competitors. This is the metric we think better represents the intensity of competition regarding resource use and spatial coexistence.

To describe coexistence in space, time and morphospace, species were grouped in pairwise symmetrical matrices, with each cell representing a species pair. We then simply filled the matrices in a binary fashion for space and time and a morphospace distance for the morphological matrix. In space and time matrices, a value of 0 represents pairs of non-coexisting/competitor species, and 1 represents pairs of coexisting/competitor species. We combined hierarchically such matrices as we increased the complexity of the competition scenario by multiplying the matrices elementwise (note this is not the same as matrix multiplication). The final product of each step was then resumed in a time series as an input to the correlation analyses described in section “*PyRate continuous analysis”* below. We described each of these time series in a scale of 0.1 million years. We devised such modular framework to be easily interchangeable as we explored new metrics of either spatial coexistence or morphospace occupation, and to be also possible to extract the intermediate steps as we described the different scenarios of competition. Supplementary figure S16 shows a “road-map” of the framework proposed here.

### Temporal and spatial coexistence

Temporal coexistence was determined by using the information of true times of speciation and extinction extracted the from each of the 50 *PyRate* replicas derived from the analyses described in section “*Initial PyRate analysis”.* The longevity of each species was defined by extracting the mean values from the posterior distributions of speciation and extinction times for each species. The number of coexisting species were measured at each 0.1-million-year discrete time interval, separately for each of the 50 replicas. We used those diversity trajectories to create 50 time series to investigate potential effects of competition as a classic scenario of diversity dependence. We are aware this likely represents a minimum estimate of the number of coexisting species because our PyRate analysis does not explicitly takes into account non-sampled species. Although this might underestimate diversity, we decided to use such metric because it will serve as base line comparison for the other time series that more explicitly take into account spatial and morphological overlap. We also notice that because the fossil record of Canidae is relatively well preserved this measure of diversity is likely to qualitatively represent the true trajectory, even if the absolute numbers are a bit off. In that respect the qualitative signal of diversity dependence is likely to be a good estimate (e.g. is there diversity dependence or not), even if the intensity of the diversity dependence is under estimated (due to missing some unsampled species). We created 373 matrices (one for each time point interval of 0.1 million years) representing the timespan of Canidae fossil record, starting from 37.2 million years ago to the present (0), for each of the 50 replicas separately. We fixed 37.2, the base of Chadronian NALMA as the origin point for all our time series. We also note that such approach considers that all species overlaps in space (or at least is spatially “perceived” by all other species), and we call this the metric “regional” coexistence.

To describe species coexistence in space in more detail, we explored a few different options that took into account properties of the fossil record of Canidae and indirectly their biology. Our intention was not to produce precise distribution maps for each species, but to try to use spatial information to describe potential scenarios of species spatial overlap and coexistence. To do so, we developed two different metrics, one very conservative (here called “Site” coexistence), which is likely to underestimate the level of species coexistence, and a more permissive one (here called “Reach distance”). Both are described shortly below but please see the supplemental material for further details.

To define spatial coexistence for the “Reach distance” approach, we used geographical information of each occurrence. From our original dataset compiled from PBDB, we extracted latitude and longitude values for all occurrences. Given the spatial and temporal resolution of the data, we deemed impossible to precisely reconstruct patterns of expanding distributions on the 0.1 million-year scale adopted to the time series of coexistence. We then proceeded to describe species coexistence adopting some simplifications. Spatial coexistence was determined using all data points found at the stratigraphic level of most occurrences, the North American Land Mammals Ages (NALMAs). Hence, two species were deemed to coexist through the whole NALMA if geographic information at the temporal resolution suggests spatial overlap.

To define spatial coexistence, we used the spatial information (latitude and longitude) of all occurrences for each species at each NALMA separately. For each pair of species, at each NALMA, we estimated the shortest distance between their occurrences and considered them to coexist if they were closer than a certain “threshold distance”. Such “threshold distance” was determined using the geographical data itself and was inspired by the potential for dispersal of each species. The overall idea was to estimate “dispersal potential” (measured as distances in km) for each species and then use this information for each species pair to determine if they were at a reachable distance (hence the name “reach”) among themselves. Thus, the “threshold distance” was different for each pair of species, measured by the sum of both species dispersal potential. Further details on this procedure can be found on the supplemental material, including Supplemental Figure S17 which describes in detail how the “reach” metric is calculated.

As a more conservative metric of species spatial coexistence (“Site” coexistence), we used an approach where only species with occurrences found in the same locality (i.e., the same fossil assemblage) are said to coexist. Using the “collection” identifier from PBDB data, we identified all occurrences of different species in the same fossil assemblage. Under this approach, species are said to coexist in space for the whole-time interval defined by the NALMA each assemblage is associated with if they are found at least once in the same fossil assemblage. Once again, we stored the spatial coexistence information in our matrices approach and multiplied those to the temporal matrices, to define species coexistence in time and space. We extracted the number of species each species coexist at a given point in time and averaged the number of coexisting species to create the time series for this competition scenario. It is important to note that no “spatial correction” such as the one applied in “reach” metric (see supplemental material) is possible, therefore, we are aware that such metric might underestimate levels of coexistence, and for that reason is considered a more conservative metric compared to the “Reach distance” method.

### Morphospace overlap

Based on the concept of competitive exclusion and ecomorphology, we hypothesize that species with very similar morphologies are expected to more strongly compete. Additionally, as the morphospace distances among species decrease and the morphospace becomes saturated, we expect the probability of speciation to decrease. To investigate such potential effect we used two distinct disparity measurements, designed to capture different aspects of morphospace occupancy through time (Ciampaglio et al., 2001; Guillerme et al., 2020).

We used the following disparity metrics: Mean Pairwise Distance (MPD) and Mean Nearest Neighbor Distance (MNND), explained below. We measured species proximity pairs using the Euclidean distances between species in a 2-dimension morphospace defined by both LD1 and body mass axis. The pairwise distance was stored in a “disparity” matrix. We latter multiply the spatiotemporal matrices described in section “*Temporal and spatial coexistence”*, either the “regional”, “reach” or “site” metric, to the matrices of ecological similarity to generate time series that describes how the average crowding of morphospace changes through time. This metric takes into account only those species that overlap in space and time. Further details on how to calculate MPD and MNND can be found in the supplemental material.

### PyRate continuous analysis

Summing up our different approaches to model different scenarios of competition, our modular framework allowed us to independently estimate metrics of purely diversity dependence competition and also incorporate more realistic mechanisms of competition. Such scenarios included either spatiotemporal aspects of coexistence or ecomorphological overlap, or even a combination of both. Time coexistence is the pivotal aspect of coexistence, incorporated across all scenarios by multiplying whatever matrices of interest by temporal coexistence. By successively increasing the complexity of competition scenarios and simultaneously analyzing intermediate steps, we expect to get a better understanding of how competition can be measured and analyzed in the fossil record. Compiling all steps of our modular framework, we ended up with 3150 time series comprising all different aspects of coexistence, competition and data completion (see supplemental material for further details).

We conducted the statistical correlation analyses between each different time series describing competition and diversification dynamics using the *PyRateContinuous* framework (Silvestro et al., 2015, 2019). *PyRateContinuous* is an implementation of the original *PyRate* software which explicitly tests if variations in speciation and extinction rates through time are associated with a given time series of a continuous variable (Silvestro et al., 2015). Under this framework, speciation and extinction rates can be associated to the time series either by a linear or exponential function, and such association can be estimated separately for different time windows. The parameter of interest is the correlation parameter, which quantifies the strength of the correlation between changes in birth-death rates and the variable of interest.

As input for this analysis, we used the species longevities extracted from our preliminary 50 *PyRate* analyses, explained in section “*Initial PyRate analysis”*. Here, we assumed an exponential correlation between changes in rates and changes in our time series in all scenarios analyzed. Given that the nature of competition might change in absolute time, due to internal or external factors, we designed our correlation analyses to be structured in different geological periods as follows: Late Eocene, Oligocene, Miocene, Pliocene and Quaternary, with boundaries at 37.2, 33.9, 23.03, 5.33 and 2.58 Ma.

We ran 30000000 MCMC iterations sampling every 1000 and discarded the first 10% as the burn- in for each analysis, separately for both across the whole timespan and for the epoch-oriented correlations. By default, the time series of interest is rescaled so that its range of values equals 1. We summarized the posterior distributions of correlation parameters of speciation (*Gl*) and extinction (*Gm*) with median values and the 95% Highest Posterior Density (HPD) credibility intervals. We also reported the baseline speciation and extinction rates, which represent the estimated speciation and extinction rates if those rates were not associated with the time series of interest (Supplementary Tables 3, 4, 5). We considered results to show strong evidence if the 95% HPD distributions did not overlap 0, however, we did not discard considerations about weaker signals of association. To test the potential association between changes in climate and changes speciation and extinction rates, we also used *PyRateContinous* to investigate if speciation and extinction dynamics were associated with changes in global temperature using the same time windows (Late Eocene, Oligocene, Miocene, Pliocene, and Quaternary) described above (see supplemental material for details). We retrieved paleotemperature data from RPANDA R package (https://doi.org/10.1111/2041-210X.12526), based on the temperature curve from Condamine et al (2013). We fit a kernel regression smooth using the function ksmooth in R to fit temperature data in our 37.2 million year time span and 0.1 million year time scale. All the R scripts used in our study are available in Appendix 1.

## Results

### Canidae diversification dynamics

Our results clearly show that speciation and extinction rates change through time, resulting in a dynamic diversification rate (Figure 1). Diversification rates are strongly positive at first, but around 30 Ma experience a marked deceleration, followed by a long phase where diversification rates were estimated to be very close to zero. Although the median diversification rate from the Late Oligocene until around the Miocene boundary is constant and positive, we should note that the 95% HPD includes zero. Through the whole Miocene, diversification rates remained very close to zero, although the median was slightly negative. There is evidence that the clade might have experienced a punctual shift in diversification around 22 Ma, and again around 13 Ma. Those shifts in diversification are not ubiquitous because the temporal placement of extinction rate shifts is highly uncertain (Figure 1B). Therefore, after the Late Eocene/Early Oligocene radiation, Canidae diversification continues to radiate followed by a modest radiation/relative stability until the Oligocene/Miocene boundary where a pulse of negative diversification started a period of gentle drop in diversity until close to the present (See Figure 2A). In the Quaternary, diversification becomes considerably negative, although the 95% HPD still includes zero.

**Figure 1:**
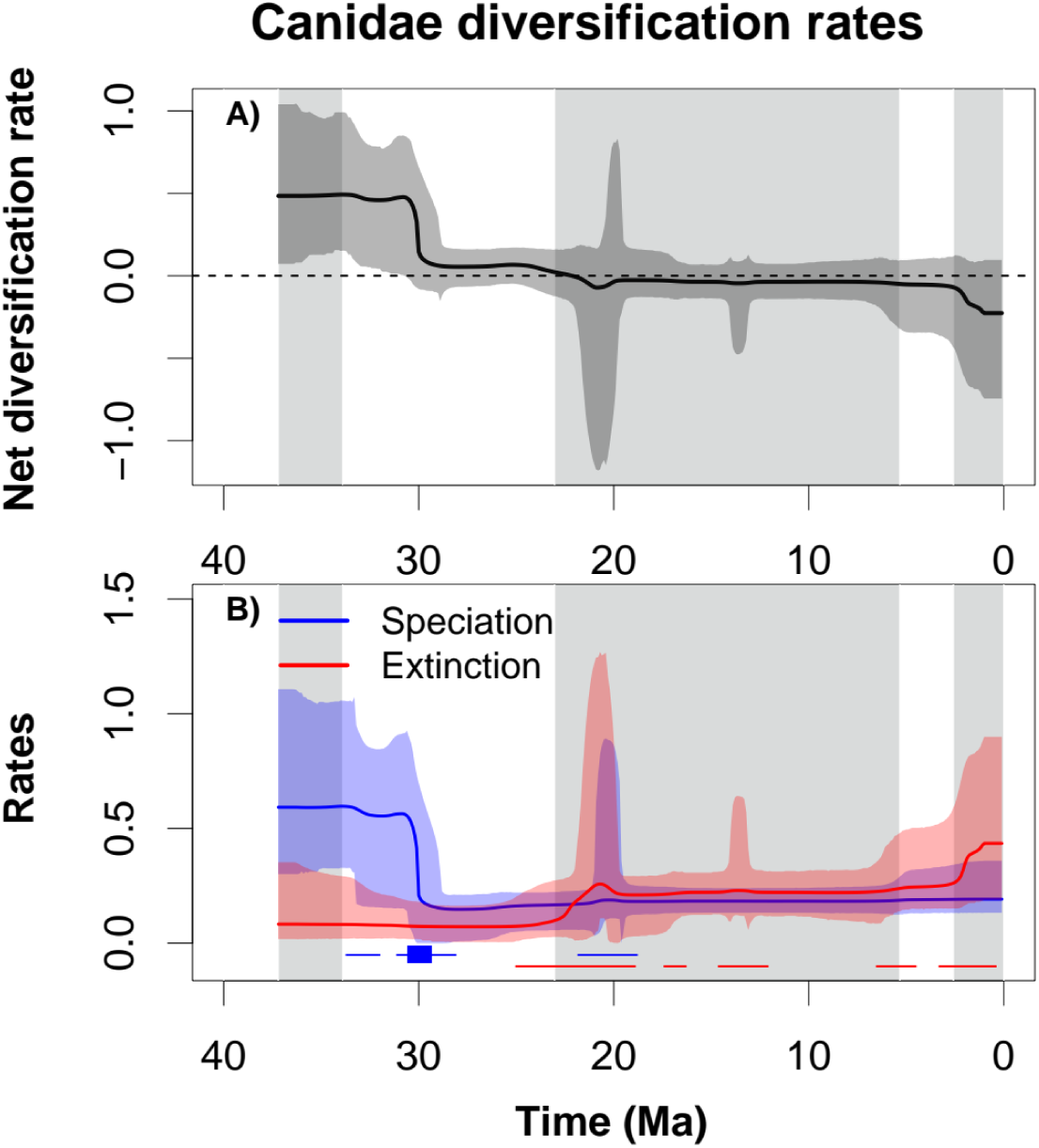
Rates through time plots showing Canidae diversification dynamics. A) Net diversification rates (speciation – extinction). B) Speciation and extinction rates. Diversification, speciation and extinction rates are expressed as the number of events per lineage per million years. Solid lines represent the median of posterior distribution, with shaded areas representing the 95% credible intervals, the Highest Posterior Density (HPD). Solid horizontal lines at the bottom represent the moments of rate shifts with BF > 2, and boxes the moments of rate shifts with BF > 6. Gray and white vertical shaded boxes represent the Cenozoic epoch boundaries at 37.2, 33.9, 23.03, 5.33 and 2.58 Ma.

**Figure 2:**
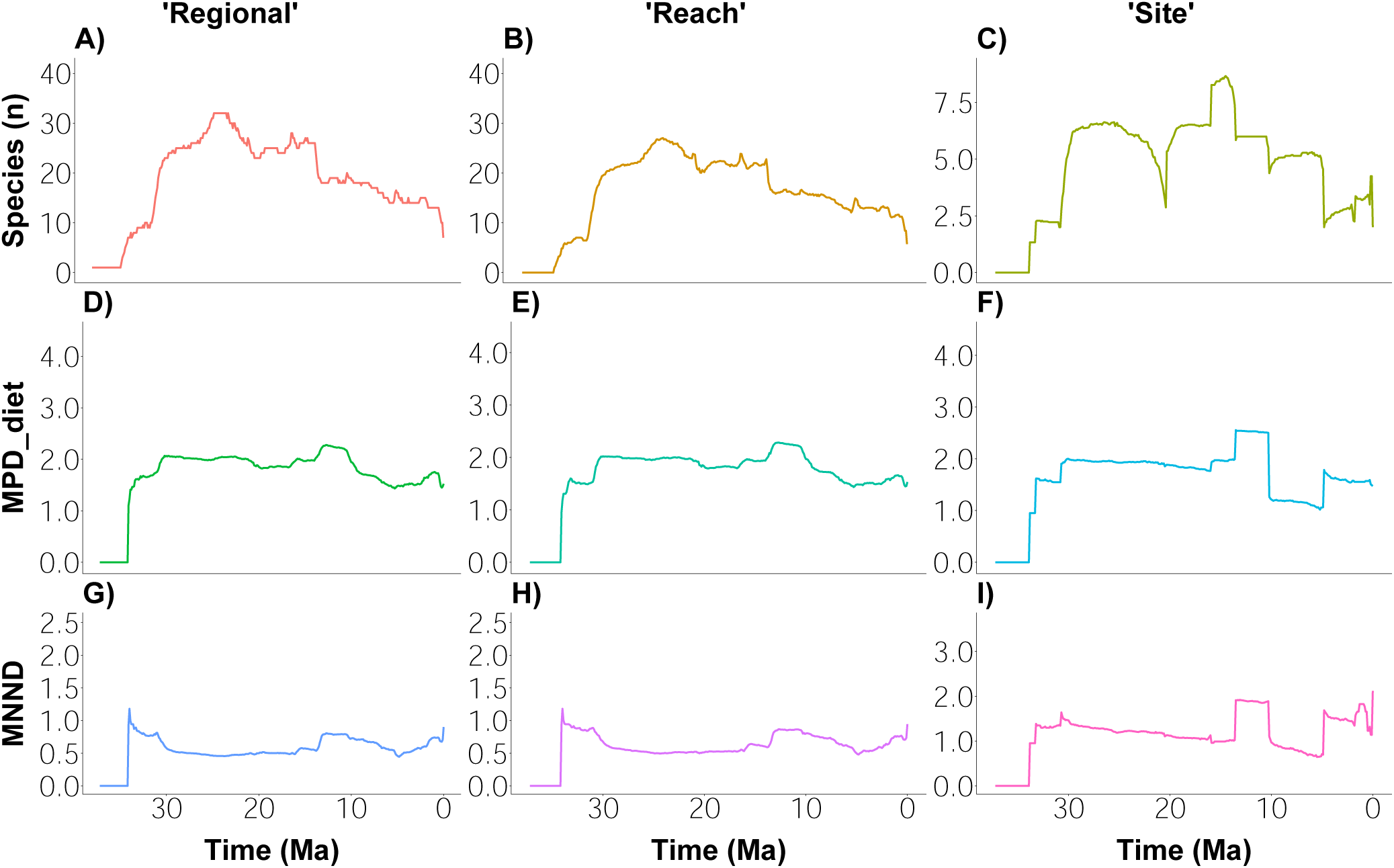
Nine different times series used as proxy for competition intensity. Solid lines represent the median value for each competition scenario, with shaded areas representing the maximum and minimum limits of each metric across the different replicas. Gray and white vertical shaded boxes represent the Cenozoic epoch age boundaries at 37.2, 33.9, 23.03, 5.33 and 2.58 Ma. Rows represent the different metrics used to measure competition intensity in morphospace, while columns represent the different approaches to measure spatial coexistence. In “regional”, species numbers are absolute, while the “reach” and “site” time series are expressed in mean number of coexisting species. MPD and MNND are dimensionless measures of Euclidean distances in the morphospace. Notice the difference of scale between metrics of spatial coexistence and MNND.

When looking at speciation and extinction rates separately (Figure 1B), we notice that the initial decrease in diversification is driven by a strong shift in speciation rates around 30 Ma, with no concomitant changes in extinction. The Bayes Factors (BF) analysis suggested strong support for this rate shift, with a BF value higher than 6 (Silvestro et al., 2019), for the combined posterior (Figure 1B), as well as for the majority of temporal replicates (Figure S3). Speciation then remained lower and constant with no strong rate shifts until the present (Figure 1B). After a period of stability through the Late Eocene and Oligocene, extinction rate increased to become slightly higher than speciation just after the Oligocene/Miocene boundary, leading to the negative net diversification. The placement of that extinction shift shows a relative temporal uncertainty, both within and between each replicate (Figure S3). Although several replicates indicate two sequential strong shifts (BF > 6) in extinction, others show a longer uncertainty in placing those shifts, which produced Bayes Factors values smaller than 6, but still higher than 2, denoting moderate support (Figure S3). Subsequent pulses in extinction might have occurred throughout the second half of Canidae history, but the support for such events is weaker (Figures 1B and S3). At the Quaternary, we detected an increase in extinction rates, although the temporal uncertainty of such shifts also leads to weaker support (2 < BF < 6) in the posterior and in many replicates. We note however, that such extinction shift towards the present is likely to be real given that it is present in virtually all replicates, with some even showing strong support (BF > 6) for such final shift (Figure S3).

### Competition scenarios and time series

Figure 2 summarizes the different time series used as our proxies for competition intensity. Through most of the Late Eocene, only one (or very few, towards the boundary depending on the replicate) species was (were) present, leading to estimates that should be taken with caution such as “zero” spatial coexistence or estimates of disparity metrics. The remaining time periods show considerably diversity and we concentrate our results and discussions on those time windows.

When looking at the simpler metric, number of sampled species within North America (Figure 2A), species diversity rapidly expands in the Oligocene, driven by high speciation rates (Figure 1B). The “regional” species richness seemed to “overshoot” around the Oligocene/Miocene boundary but remained relatively stable through most of the Oligocene and Miocene until about 15 Million years ago. Diversity then experienced a quick drop, and then a steady and subtle decline throughout the Miocene and Pliocene. Very close to the present, “regional” species richness abruptly drops to reach current number of species. The “reach” spatial coexistence scenario (Figure 2B) closely mirrors the “regional” diversity pattern, although with slightly lower values (see also Figure S8 showing the correlation of those two time-series).

The “site” scenario displays both similarities and striking differences to the other metrics. As with the other metrics it suggests a general trend of “rise and fall” in the number of coexisting species. The phase of expanding diversity suggests a very abrupt rise in the average number of coexisting species at the Early Oligocene. The remainder of the period, and part of the Miocene, presents a roughly constant number of coexisting species, punctuated by two events: one showing a dramatic drop and immediate recover in the average number of coexisting species (around 20 Ma), and another showing a momentary increase in the average number of coexisting species (around 13 Ma). Such marked decline and subsequent rise in diversity are not seen in the other diversity time series (Figure 2A, B). Through the rest of the Miocene it follows two short periods of roughly constant numbers of coexisting species interrupted by a stepwise drop. Hence, the most striking differences between the “site” time series and the other measures of spatial coexistence are: i) there are more stepwise changes in diversity throughout time, and ii) the gentle increase in the number of coexisting species throughout most of the Pliocene and Pleistocene (only seen in “site” coexistence).

The different time series depicting changes in Mean Pairwise Distances (MPD) are mostly concordant (Figure 2D, E, F) and show in fact a very strong correlation (Figure S8). Contrary to our expectations, MPD did not change much throughout most of Canidae history. Apart from the first abrupt increase in the Eocene/Oligocene boundary, which is likely an artifact resulting from estimating distances at very low richness in the Eocene, the MPD values remained roughly constant throughout the whole time. The main exceptions were two stepwise changes at the Late Miocene when we used the “site” spatial coexistence metric (both temporally concordant with the stepwise changes seen in “site” diversity). During the Canidae radiation (Late Eocene/Early Oligocene) the MPD values, if anything, increases instead of decreasing as expected.

Mean Nearest Neighbor Distances (MNND), showed temporal trends that better corresponded to our expectations of increasing competition, at least when considering the “regional” and “reach” approaches. During the expanding phase of diversity, MNND decreases in both “regional” and “reach” coexistence (Figure 2G, H). In other words, as more species are added in the morphospace, the distance between a species and its nearest competitor becomes smaller, even at the face of an increase in total morphospace for the same time period (Figure S7). Through the Miocene, MNND values remained constantly low for the “regional” and “reach” approaches (Figure 2G, H), but showed an increase during the Pliocene and Quaternary. Such increase in MNND was accompanied by decreases in species diversity (Figure 2A, B), and an increase in total morphospace area (Figure S7), suggesting a scenario of less intense competition among Canidae species, where species were more sparsely distributed in the morpho and geographical space. However, these patterns were only seen for the “regional” and “reach” methods. The “site” MNND time series did not show the steady decrease seen in Canidae radiation during the Oligocene (Figure 2I), nor the steady rise from the Miocene/Pliocene boundary up to the present. Instead, it showed marked stepwise shifts occurring from the second half of the Miocene up to the present. Those seemed also temporally congruent with the stepwise changes seen for the “site” diversity and MPD, which suggests that changes in “site” diversity might be driving those stepwise changes when we measure disparity at the “site” spatial scale.

### Association between macroevolutionary dynamics and the intensity of competition

When considering only the number of coexisting species (Figure 2A, B, C), all metrics used suggested strong evidence for negative correlation between changes in speciation rate and changes in the diversity/coexistence time series during the Oligocene (Figure 3). For this time interval, the 95% Highest Posterior Density (HPD) interval did not overlap with 0. Such pattern of negative correlation is only present during the phase of diversity expansion (Oligocene), whereas in other time periods, where diversity is roughly constant or decreasing, no evidence for correlation was detected (Figure 3A, B, C). MPD analyses did not show any association between any of the different time series and speciation rates (Figure 3D, E, F). We decided to still report those results because this lack of association can be seen as some kind of “positive control” for our statistical analysis (at least when compared to the MNND analysis), and because it helped us to get some interesting biological insights. Given the roughly constant value of MPD (Figure 2D, E, F), this is not surprising, but we suspect that such lack of association might have also been affected by a concomitant change in the total morphospace area (Figure S7) and some kind of dilution effect when using the MPD metric because we are averaging out distances of species that are really far apart in the morphospace which are unlikely to interfere with each other. During the Oligocene, as the number of species increased, the total morphospace area increased allowing the mean distance between species pairs to either remain constant or even slightly increase.

**Figure 3:**
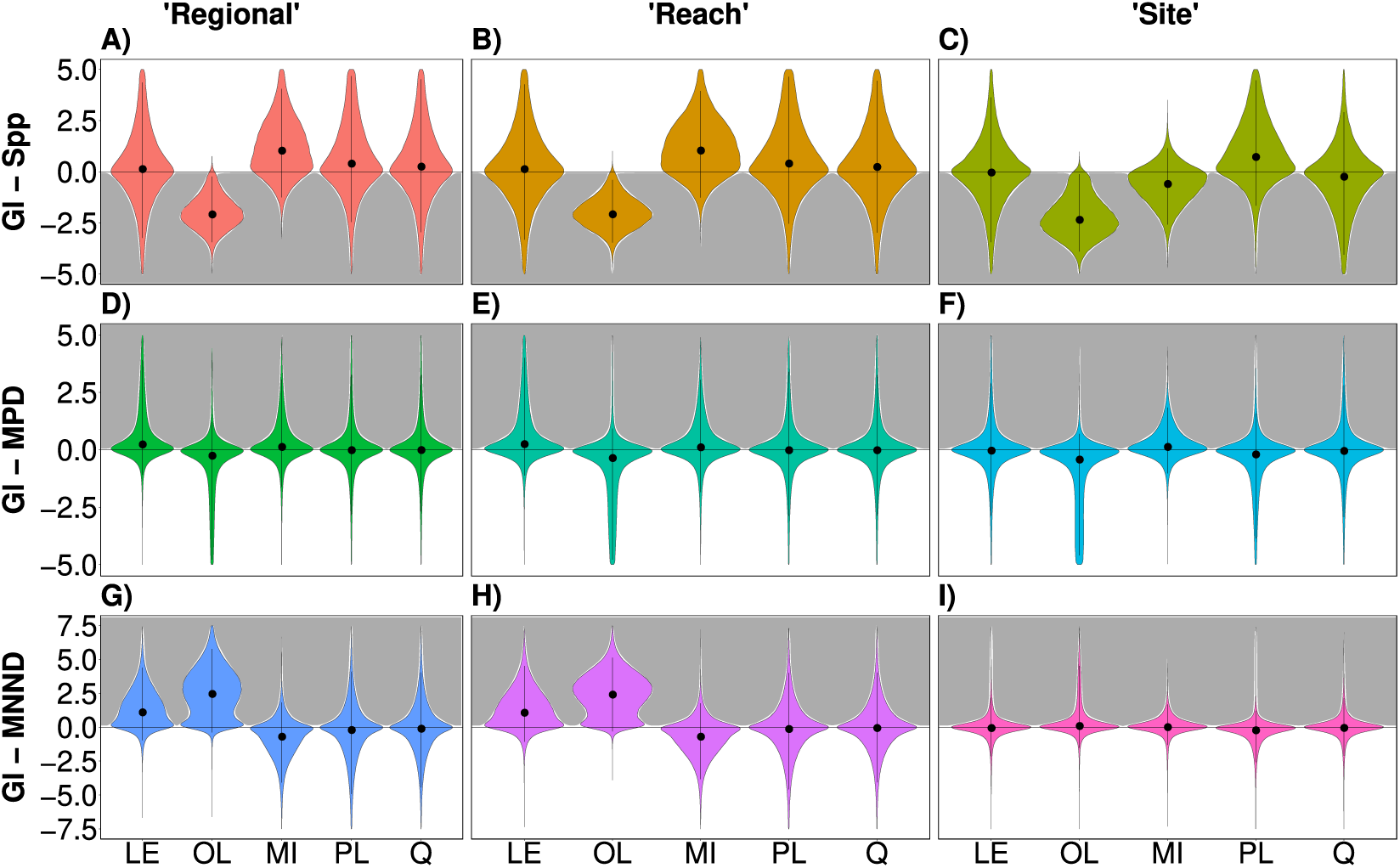
Posterior distributions of speciation correlation parameters (Gl) in discrete time windows. LE: Late Eocene, OL: Oligocene, MI: Miocene, PL: Pliocene, Q: Quaternary. Correlation parameters are dimensionless and quantify the amount of change in rates associated with the change in the time series of interest. Rows, columns and color-coding are the same as in Figure 2. Gray shaded areas highlight our expectations for the values of the parameter if speciation rate responded accordingly to what would be expected if competition was a relevant factor, with positive or negative values depending on the metric. The vertical black lines within each violin plot represents the 95% HPD and the dot represent the median of each combined posterior distribution.

During the Late Eocene and Oligocene time windows, the *Gl* 95% HPD describing the potential association between MNND and speciation rate showed evidence for a positive correlation for the “regional” and “reach” coexistence scenarios. Although this positive correlation might seem at first counter-intuitive (compared to the negative correlation seen for the number of species) we should note that MNND measures the minimum distance between two neighbor species. Therefore, the positive correlation means that as MNND decreases species becomes closer to each other, and we expect intensity of competition to increase, leading to deceleration in speciation. During the Late Eocene, the rapid expansion of diversity from one to only a few species (Figure 2A) leads to a rapid increase in MNND from 0 to about 1 (Figure 2G). Given that such dynamics was measured in a very small interval with very few taxa, we suggest that such correlation must be viewed with caution.

Although we see evidence for correlation between changes in competition intensity and speciation rate during the Oligocene (when using “regional” and “reach” MNND), we note that the 95% HPD do cross the value of zero, weakening the evidence for this correlation (Table S3). In both cases (“regional” and “reach” MNND) we see a rather bimodal pattern, where the major mode is clearly offset from the value of zero, but the minor mode is very close to zero (Figure 3G, H, table S3). When we look at the posterior distributions of each replicated dataset independently (Figures S9 and S10), we note that the median of every dataset is offset from the value of zero, but some (less than half) replicates have their 95% HPD crossing the value of zero. This indicates that this uncertainty seen in the combined posterior distribution mostly comes from differences between replicates even though there are some uncertainty within some replicates as well. We detect no evidence of correlation between “site” MNND and speciation rate, a result somewhat expected given the trajectories observed in these individual time series (Figures 2I and 3I).

The results for the extinction dynamics showed no strong correlation between the different proxies for competition and extinction rates (Figure 4). The *Gm* posterior distributions across all competition scenarios are usually centered at or very close to zero, and in few time windows where this was not the case, the 95% HPD crossed zero, and showed a correlation in the opposite direction as expected by a competition scenario (Figure 4). In the only time interval (Quaternary) where we see some reasonable evidence for an association, the evidence was weak and in favor of a pattern that is the inverse of what would be expected if competition was relevant. This unexpected association was detected for the “regional” and “reach” analyses of species coexistence (Figure 4A, B), and their corresponding MNND analyses at the Quaternary time window (Figure 4G, H). Here, the 95% HPD is off centered towards positive values for the “regional” and “reach” coexistence analyses and towards negative values for their corresponding MNND analyses.

**Figure 4:**
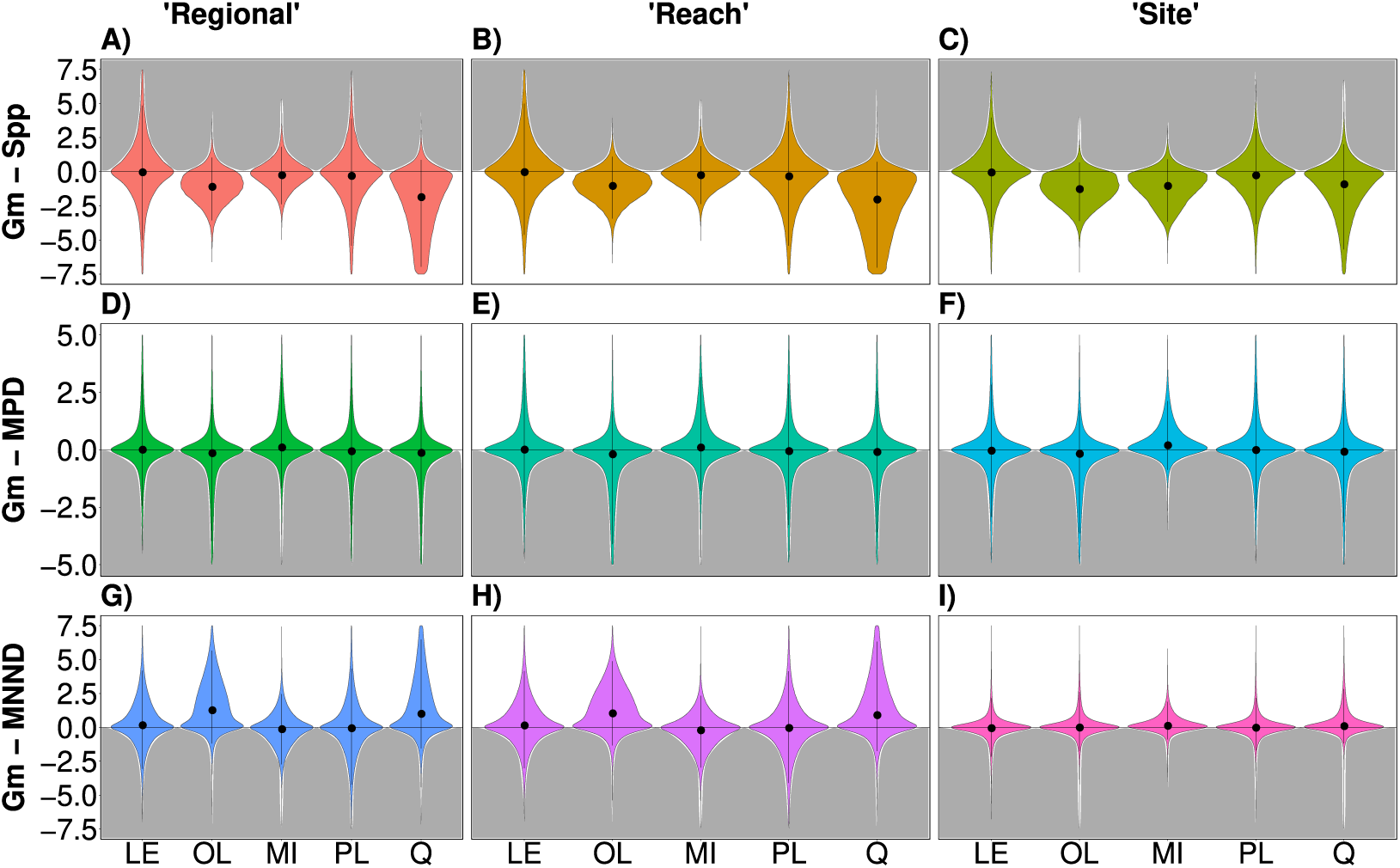
Posterior distributions of extinction correlation parameters (Gm) in discrete time windows. **LE:** Late Eocene, OL: Oligocene, MI: Miocene, PL: Pliocene, Q: Quaternary. Correlation parameters are dimensionless and quantify the amount of change in rates associated with the change in the time series of interest. Rows, columns and color-coding are the same as in Figure 2. Gray shaded areas highlight our expectations for the values of the parameter if extinction rate responded accordingly to what would be expected if competition was a relevant factor, with positive or negative values depending on the metric. The vertical black lines within each violin plot represents the 95% HPD and the dot represent the median of each combined posterior distribution.

Changes in global temperature are not associated with changes in speciation nor in extinction when we consider the same time intervals considered before (Figure 5). When considering the Cenozoic as a whole we see evidence that changes in temperature are associated with changes in extinction dynamics (Figure S15).

**Figure 5:**
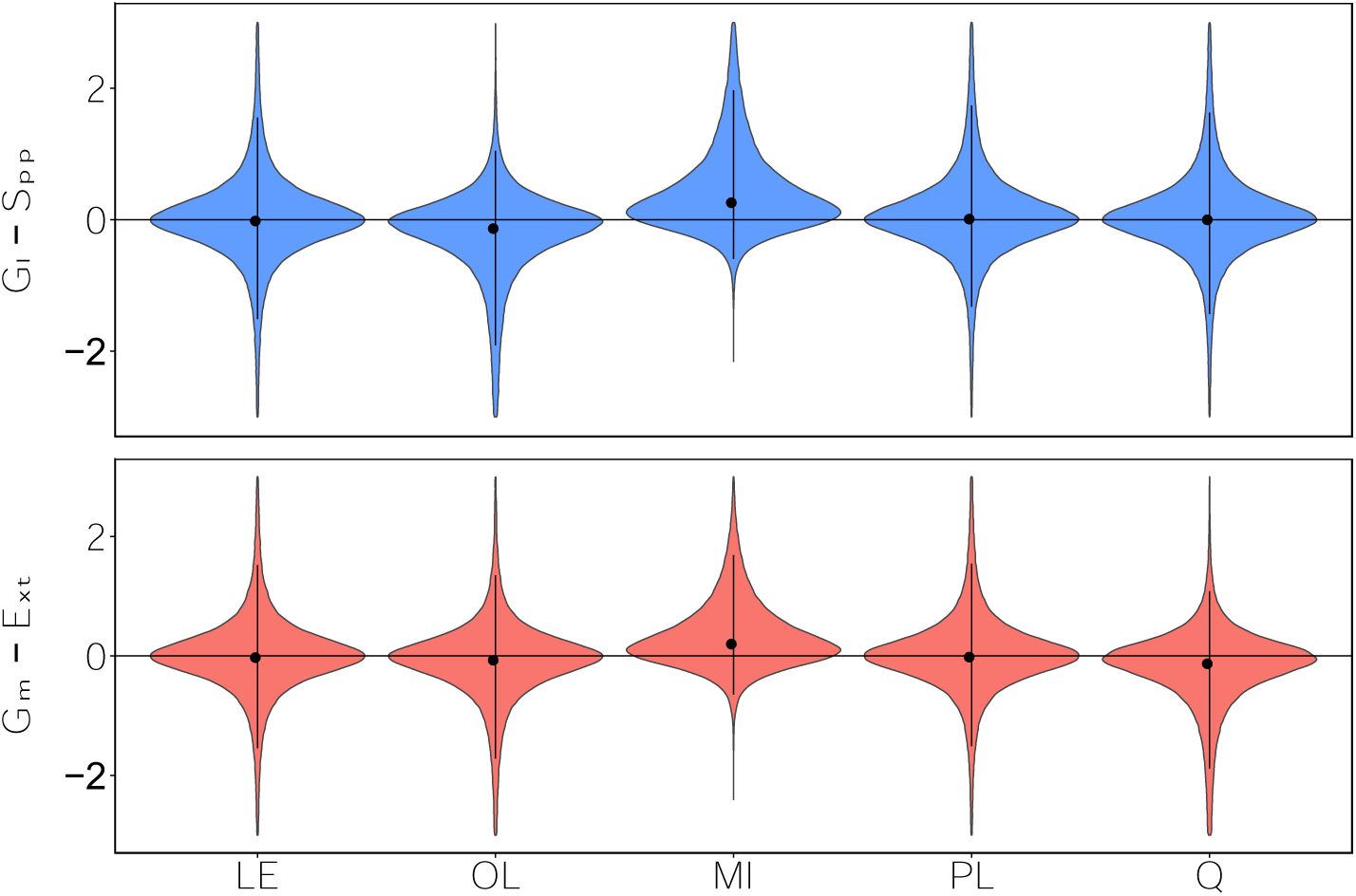
Posterior distributions of speciation (Gl) and extinction correlation parameters (Gm) in discrete time windows. LE: Late Eocene, OL: Oligocene, MI: Miocene, PL: Pliocene, Q: Quaternary. Correlation parameters quantify the amount of change in rates associated with the change in the time series of interest. Rows, columns and color-coding are the same as in Figure 1B. The vertical black lines within each violin plot represents the 95% HPD and the dot represent the median of each combined posterior distribution.

## Discussion

Studying the effect of species interactions in deep time is challenging because we cannot directly observe the outcome of such interactions (Ezard et al., 2016). Although diversity dependence has been previously described and used as a proxy for interspecific competition in several lineages (Condamine et al., 2019; Foote et al., 2018), including Canidae (Pires et al., 2017; Silvestro et al., 2015), the notion that resource competition is the underlying mechanism remains an indirect interpretation due to the nature of deep time studies. In our study, we did not fully overcome this historical barrier, but we employed an approach that more explicitly inferred resource competition by measuring how the amount of niche and spatial overlap changed over time and examining whether these changes were associated with variations in diversification dynamics. By doing this we were also able to compare our inferences using different proxies for interspecific competition, and hence get a better insight on the potential role of interspecific competition.

The initial expansion of Canidae diversity is mostly driven by the appearance of hypo and mesocarnivorous morphologies (Figures S12 and S13), while the hypercarnivorous niche was mostly occupied by other clades such as Hyaenodonts and Nimravids (Van Valkenburgh, 1999). Right after the initial occupation of the morphospace, when diversity started to rise, many Canidae hypercarnivorous forms start to appear roughly at the same time around 30 Ma (Figures S12 and S13; see also Slater 2015). The rapid expansion of early hypercarnivorous Canidae forms is hypothesized to have actively influenced the demise of Hyaenodonts (Van Valkenburgh, 1999), with recent studies demonstrating such statistical evidence for such displacement (Pires et al., 2017). In the context of competition imposed by species within the family Canidae, this rapid filling of the morphospace translates into a progressive drop in the MNND metric (Figure 2G, H), a concomitant drop in speciation (Figure 1B), and hence a more direct evidence for a role of resource competition on driving the speciation dynamics during the Oligocene (Figure 3G, H). Our results suggest that when taking into account temporal and morphological overlap, we see strong evidence for interspecific competition suppressing speciation rates at least in two out of the three spatial scales used here (Figure 3). Our diversity dependence results are also consistent with previous analyses that evaluated such models at the continental regional scale (Silvestro et al., 2015; Pires et al., 2017). Interestingly, our new results from the “reach” and “site” metrics, designed to explicitly incorporate spatial coexistence at lower spatial levels, also showed strong evidence for diversity dependence (figure 3 A to C; but see supplemental material for brief discussion on limitations of diversity-dependent inference). Taken together, these results suggest that interspecific competition can be captured at different spatial scales, and that irrespective of the spatial scale, it seems to operate only through the speciation dynamics as proposed by Pires et al (2017), and only at certain time windows as suggested here.

The most unexpected result was the absence of association between changes in MNND and speciation rates when we considered spatial coexistence at the “site” metric (Figure 3I). This was somewhat surprising given that MNND showed evidence for association with speciation dynamics for the other metrics of spatial coexistence (Figure 3G, H), and diversity dependence at the site scale (Figure 3C). It is worth noticing that through the Oligocene, the moment with a significant strong shift in speciation rates, we see a steady drop in MNND values the “regional” and “reach” scenarios, while in “site” scenario, MNND remains roughly constant (Figure2; with a sharp peak around 30 Ma), and at a considerable higher value than the others (see Figure S14). Given that MNND metric at the site measures the average the average distance in morphospace that each species coexisted and interacted with locally, the lack of an association between MNND at the site level and speciation dynamics could be due to either a biological or sampling effect.

Limiting similarity might be so strong and effective that it results in local fossil assemblages where species are more dissimilar than what the regional pool of species would suggest . Similar species might come into contact, but they would not coexist long enough to be recorded in the fossil record at the site scale. Extant canids avoid sharing the habitat when competition is high (Johnson et al., 1996), either by dividing habitat use or food resource, or by establishing dominance hierarchies. If we assume that extinct canids also had such interference/avoidance at the local scale, we could expect that this metric would detect such dynamic interference more easily. It is also possible that local coexistence leads to character displacement (Brown and Wilson 1956; Grant and Grant 2006), a phenomenon that has been also detected in the fossil record (Schindel and Gould 1977). In the fossil record, only species different enough (either by species sorting and/or character displacement) would survive long enough to be found at the same locality. The other metrics, by measuring overlap at a larger spatial scale (“reach“), or by assuming that all species “sense” each other in space (“regional“), would reflect an overlap at a larger spatial scale, even if at the local scale this coexistence is ephemeral. Therefore, using different spatial scales might help us to capture the different aspects of the coexistence dynamics, and we should not expect them to always yield similar results. Another possibility is that the lack of association between changes in MNND at the site and changes in speciation rate is due to the fossil record being incomplete. The fossil record usually improves in quality at higher taxonomic and spatial scales, and the “site” metric might be particularly affected by the record incompleteness. If a pair of species do not fossilize at the same location, we would not infer their coexistence using this metric. Therefore, even though it is tempting and reasonable to interpret the differences between the site and other spatial levels, we have to keep in mind this potential limitation.

Contrary to the initial theoretical propositions of interspecific competition based on diversity dependence (Sepkoski, 1996), we see that changes in interspecific competition intensity (either measured by changes in diversity or changes in the morphological average distances between spatially coexisting species) did not result in changes in the extinction dynamics. We note that at the moment Canidae family was experiencing its initial radiation, extinction remained constant (Figure 1B), even in a progressively more crowded morphospace, at least as measured by changes in MNND (Figure 2G, H). We suggest that the apparent irrelevance of interspecific competition on extinction dynamics during the radiation phase might be related to the speciation mechanism itself, and to some extent to “self-regulating” mechanism that operate at the clade level. If ecological speciation is relatively common (Nosil, 2012), and new species overlap, even partially, in space and time with their progenitors, then it is possible that speciation produces only species that are already ecologically distinct, limiting the effect that competition might have on the survival of a given species once they are fully formed. This reasoning applies to “fully fledged species” (instead of incipient or ephemeral species *sensu* Rosenblum et al 2012, Dynesius & Jansson, 2013), which is what we are likely to be measuring when using the fossil record.

This is not to say that interspecific competition will never affect the extinction dynamics, but the phylogenetic distance and geographical isolation might determine the pool of species that most likely interfere with the extinction dynamics of a given clade. Previous evidence in favor of diversity-dependent extinction was typically detected among clades and not within clades (Pires et al., 2017; Silvestro et al., 2015). Moreover, in those cases most of the interspecific competition among species in different clades involved a clade that migrated from another continent (Pires et al., 2017; Silvestro et al., 2015). In the particular case of Canidae, Silvestro et a (2015) and Pires et al (2017) have shown that North American Canidae family, and its sub-families, extinction dynamics is strongly associated with changes in Felidae diversity after their arrival from Eurasia sometime during the Miocene. Future work might implement the approach proposed here to measure the intensity of competition among clades and hence better evaluate the potential effect of clade competition on extinction dynamics.

Climate change likely shaped much of mammalian diversification in the last 20 Million years (Janis, 1993), driven mostly by the transition to the “Icehouse Earth” in Late Cenozoic (Smith & Pickering, 2003). Cooling climates driven by reconfiguration of oceanic currents and formation of ice caps lead to the turnover of vegetational habitats across the world, with increasing diversity of open grasslands in North America, at the demise of denser forests of the Early Cenozoic (Strömberg, 2002, 2005). The transition to Icehouse climates started at the Eocene-Oligocene boundary has been proposed to be a relevant force in driving Eurasian mammal diversification dynamics (Hooker et al 2004). Pires et al (2017), as well as Balisi & Van Valkenburgh (2020) found an overall (across the whole Cenozoic) negative association between changes in Canidae extinction rates and temperature. We also detect such pattern when considering the whole-time interval (Supplemental figure 15), but no effect when testing the association at different time windows, neither for speciation nor extinction rates. We interpret this discrepancy between analysis using different short time windows vs. a single time window as evidence that the long-term changes in temperature rather than smaller changes happening at different time windows, were relevant. In the case of canids, a long-term decrease in temperature seems associated with an increase in extinction rate which suggests the inability of Canidae species to adapt to colder climate. The negative statistical association might also simply represent an “correlation” effect and not causality (Liow et al 2015; Hannisdal & Liow 2018) and future work might try to disentangle such scenarios.

Initial debates on whether biotic or abiotic factors might control biodiversity in deep time have played one factor against the other resulting in somewhat artificially dichotomic views on deep time controls of biodiversity. Although there is a growing body of evidence suggesting a role for biotic factors (Foote et al., 2018; Liow et al., 2015), and it is well established among paleontologists that, at deep time, abiotic factors are undoubtedly relevant (Barnosky, 2001; Erwin, 2009; Hannisdal & Peters, 2011; Jaramillo et al., 2006; Mayhew et al., 2012; Peters, 2008; Welker et al., 2015), it has been recognized that biotic and abiotic factors might interact to produce changes in biodiversity (e.g. Ezard et al., 2011).

Our temporal framework allowed us to suggest that interspecific competition (either measured as diversity dependence or more interestingly, by the amount of temporal-spatial-niche overlap) is not an omnipresent mechanism throughout entire Canidae radiation. By incorporating spatial and ecological overlap more explicitly, and discretizing different time windows, we have demonstrated that interspecific competition only took part in suppressing speciation rates in moments when the diversity was increasing at the initial radiation. We propose that the relevance of biotic and abiotic factors might in fact change according to the lineage’s “ontogeny”, and recent work has also found evidence for a similarly more complex scenario in gastropods (Neubauer et al 2022). For Canidae, we propose that biotic factors driven by interspecific competition might be more relevant at the initial stages of lineages, while external factors such as climate change might be more relevant as a long term effect through. Hence, although it is clear that in the “ecological theater and evolutionary play” (Hutchinson, 1965) we should not have a monologue of either the Red Queen^1^ or the Court Jester, it is likely that the specific leading roles changes depending on the play act.

## Funding and Acknowledgements

R.P.G. thanks CAPES for a master’s program fellowship, L.P thanks FAPESP for a posdoc fellowship (# 2022/03664-6), and T.B.Q thank FAPESP for financial support (grant # 2021/06780-4). We would like to thank Mathias Pires, Daniele Silvestro, and all members of the LabMeMe for providing valuable feedback. We would also like to thank all scientists who have contributed data to the Paleobiology Data Base (PBDB). This is PBDB publication #xxxx.

## Footnotes

1 We note that even though in his original formulation of the Red Queen hypothesis, Van Valen (1973) gave special attention to biotic factors, he considered changes in a deteriorating environment to be caused by both biotic and abiotic factors. More recently, researchers have restricted the term Red Queen to biotic factors and Court Jester to abiotic factors.

